# Visualizing and identifying selfish bacteria: a hunting guide

**DOI:** 10.1101/2023.05.10.540300

**Authors:** G. Reintjes, G. Giljan, B. M. Fuchs, C. Arnosti, R. Amann

## Abstract

Polysaccharides are dominant components of plant biomass, whose degradation is typically mediated by heterotrophic bacteria. These bacteria use extracellular enzymes to hydrolyze polysaccharides to oligosaccharides that are then also available to other bacteria. Recently, a new mechanism of polysaccharide processing – ‘selfish’ uptake – has been recognized, initially among gut-derived bacteria. In ‘selfish’ uptake, polysaccharides are bound at the outer membrane, partially hydrolyzed, and transported into the periplasmic space without loss of hydrolysis products, thus limiting the availability of smaller sugars to the surrounding environment. Selfish uptake is widespread in environments ranging from the ocean’s cool, oxygen-rich, organic carbon-poor waters to the warm, carbon-rich, anoxic environment of the human gut. We provide a detailed guide of how to hunt for selfish bacteria, including how to rapidly visualize selfish uptake in complex bacterial communities, identify selfish organisms, and distinguish the activity of selfish organisms from other members of the community.

## Introduction

Polysaccharides constitute the largest pool of metabolically accessible organic carbon in the biosphere ^1^. Their primary sources are phototrophic organisms of the terrestrial and marine environment, which produce polysaccharides as structural complexes and as storage compounds ^2-4^. Polysaccharides account for about half of the living biomass of phytoplankton ^3^ and terrestrial plants ^5^ and represent a major fraction of the immense reservoir of detrital organic matter in soils ^6^, sediments ^7^, and seawater ^8^. The cycling of polysaccharide-derived material thus is critical for processes and issues ranging from the global flux of carbon to human ^9-11^ and animal ^12, 13^ nutrition.

Polysaccharide degradation, transformation, and remineralization is mainly performed by bacteria, which are abundant in the environment ^14^ and in the digestive tracts of animals ^15^. Degradation of polysaccharides is challenging for bacteria because polysaccharides are structurally complex ^16^, containing different monosaccharides connected by a wide range of glycosidic linkages ^16, 5^. Since these monosaccharides can be linked together via any of five or six positions, the structural complexity of polysaccharides far outpaces that of other biopolymers, such as proteins. Thus, correspondingly complex enzymatic systems are required for polysaccharide degradation ^17^.

Initial enzymatic hydrolysis of polysaccharides by bacteria occurs outside the cell due to the large size of polysaccharides. This extracellular hydrolytic step produces lower molecular weight products that can be released into the surrounding environment and may be available for uptake by organisms that did not produce the extracellular enzymes (Fig. 1 A, C, E) ^18-20^. This potential loss of hydrolysis products constitutes a complication for extracellular enzyme-producing bacteria, which need to obtain sufficient hydrolysate as a return on their investment in hydrolytic enzymes. Recently, however, a distinctly different mechanism of polysaccharide processing – ‘selfish’ uptake – has been recognized (Fig. 1 A, B, D), initially in gut bacteria. ‘Selfish’ bacteria ^21^ bind, partially hydrolyze, and transport polysaccharides into the cell, releasing little to no low molecular weight hydrolysis products to the surrounding environment, thereby ensuring a return on their enzymatic investment.

**Fig. 1.**
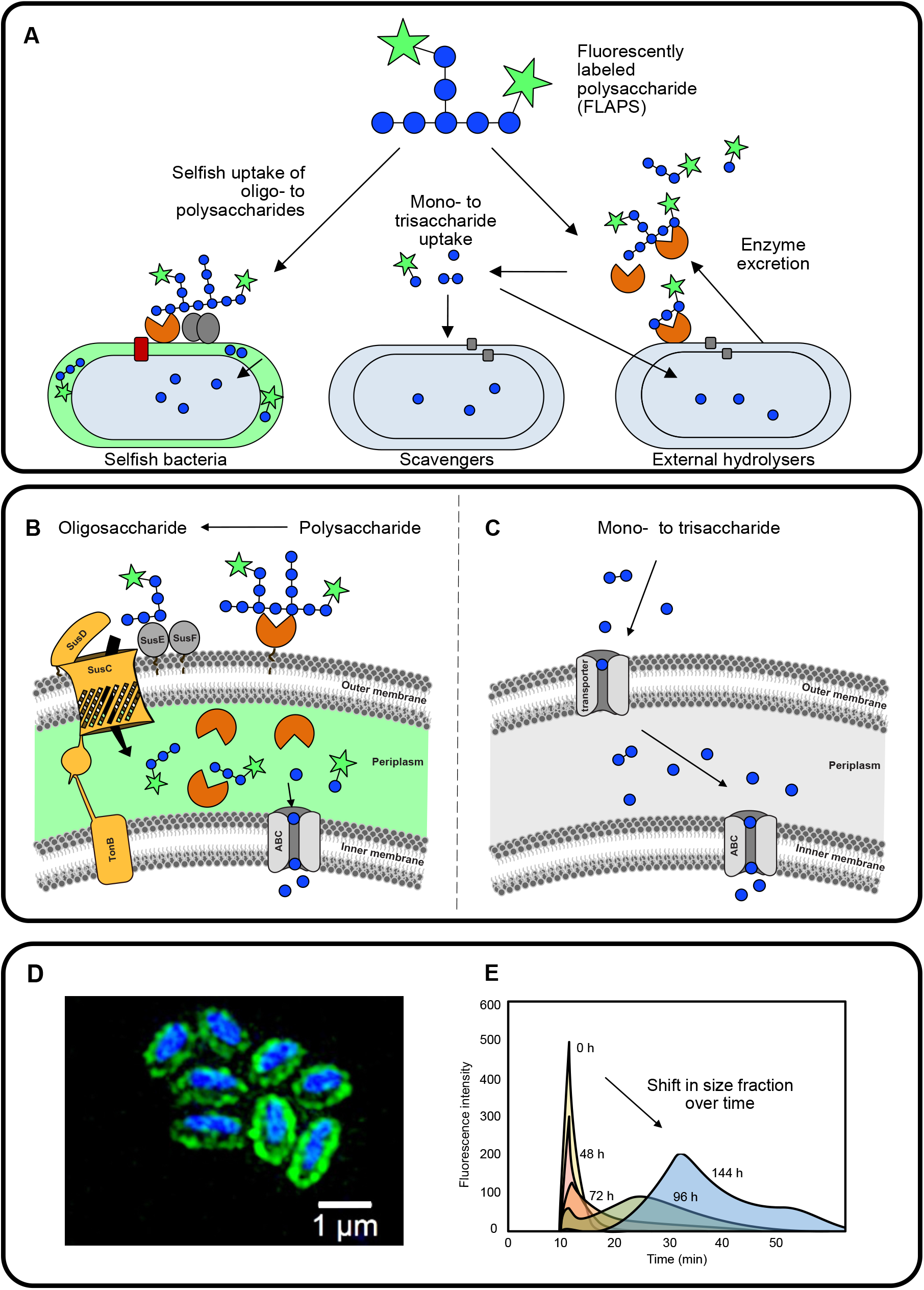
Heterotrophic utilization of polysaccharides as shown with a fluorescently labeled polysaccharide (FLAPS). **(A)** Schematic overview of the two known polysaccharide utilization mechanisms – selfish uptake and extracellular hydrolysis with subsequent uptake of monosaccharides and small oligosaccharides. **(B)** Conceptual schematic of selfish FLAPS uptake into the periplasm of a cell where it is further hydrolyzed to monosaccharides that are transported through the inner membrane into the cell. (Adapted from Hehemann et al., 2019^13^) **(C)** Conceptual schematic of enzymatic extracellular FLAPS hydrolysis into monosaccharides that are transported into the cell. (Adapted from Arnosti et al., 2018^19^) **(D)** Microscopic visualization of selfish FLAPS uptake and accumulation in the cells. **(E)** Gel permeation **c**hromatogram showing systematic changes in molecular weight of FLAPS with incubation time (0 – 144 h) (Adapted from Arnosti, 2003^30^).

The cost of enzyme production, and the complexity of enzymatic systems required to deconstruct many polysaccharides, thus, may be balanced in different ways. Selfish uptake likely requires high energetic investment to express many enzymes but is characterized by little loss of hydrolysis products ^21^. External hydrolysis potentially leads to the loss of low molecular weight hydrolysis products to other organisms, but might be coordinated among bacteria (e.g., via quorum sensing ^22^), such that enzyme production and hydrolysate uptake can be optimized within a community. Initial assessments of the prevalence of selfish uptake and external hydrolysis in the ocean suggest that strategies of substrate processing change with location, as well as with the nature and abundance of substrates ^20, 23, 24^. In particular, selfish uptake may pay off, particularly in cases where competition for a specific substrate is very high, as well as in cases where the abundance of a complex substrate is low, such that a return on investment in complex enzymatic systems needs to be guaranteed ^25^.

In sum, ‘selfish uptake’ is prevalent among organisms found in the anoxic, organic-carbon-rich gut environment ^26^, and also in the oxygenated organic carbon-poor waters of the surface ocean ^27^. The recent discovery that selfish bacteria are also abundant throughout the oceanic water column and take up substrates that are not hydrolyzed externally demonstrates that standard methods to determine microbial activities may be overlooking important organisms ^28^. In short, given their presence in these distinctly different environments, selfish bacteria are likely to be found in many other natural environments, including sediments, soils, and digestive tracts of a wider range of organisms. Therefore, detecting the presence and activities of selfish bacteria is central to our efforts to understand carbon cycling, animal nutrition, and the microbial ecology of a wide range of environments. Fortunately, detecting the presence of selfish bacteria and selfish activity experimentally is a straightforward process.

Identifying the presence of selfish bacteria also opens the door to further focused investigations, starting with the taxonomic identification of selfish bacteria and extending to flow cytometric methods that enable the physical separation of these bacteria and further analysis of their physiology, biochemistry, and activity. We present an example from the North Sea demonstrating how hunting for selfish bacteria can yield further information about community activities, identities, and carbon flow in a natural system. These data were initially presented in Giljan *et al*., (2022^29^); here, we present in detail results that were not discussed at length in that manuscript. We also discuss additional insights from human gut microorganisms ^13^. The approach we used could easily be applied to studies in fields ranging from animal nutrition to terrestrial and aquatic investigations of the ecology of microbial communities and the pathways of carbon degradation that they catalyze in natural environments. Overlooking selfish bacteria and their activities in any environment means that we are overlooking important organisms, as well as pathways of material flow and energy transfer. Here, we present in detail the methods required to reveal their presence.

## Results

### A rapid workflow to detect active polysaccharide utilizers

Fluorescently labeled polysaccharide (FLAPS) incubation experiments are, at present, one of few methods to provide insights into the mechanisms of polysaccharide processing – extracellular hydrolysis and selfish uptake – with the possibility to link the function to the identity of specific bacteria: to date, selfish uptake cannot be detected solely via ‘omic analyses ^26^. Here, we present a simple approach to detect bacteria – in pure cultures and complex environments – that are actively taking up polysaccharides through a selfish mechanism (Fig. 2). In brief, the FLAPS of interest is added to a liquid sample or culture medium, incubated, and subsamples are periodically collected and filtered. Selfish substrate accumulation in the periplasm can be visualized with a standard epifluorescence microscope after initial DNA staining (e.g., DAPI) through co-localization of the FLAPS signal and the nucleic acid counter stain. FLAPS signals without a nucleic acid counterstain should be excluded as background noise. This simple and straightforward approach allows simultaneous quantification of total and selfish cells, and answers the first key question: are selfish bacteria that use this specific polysaccharide active in my sample? This experimental setup also permits further investigations: the same filter can be used to analyze bacterial community composition by 16S rRNA amplicon sequencing (Supplementary Fig. S1D). Moreover, collecting the filtrate allows measurement of extracellular hydrolysis rates, using gel permeation chromatography and fluorescence detection (Fig. 1E, Supplementary Fig. S1A) ^30^.

**Fig. 2.**
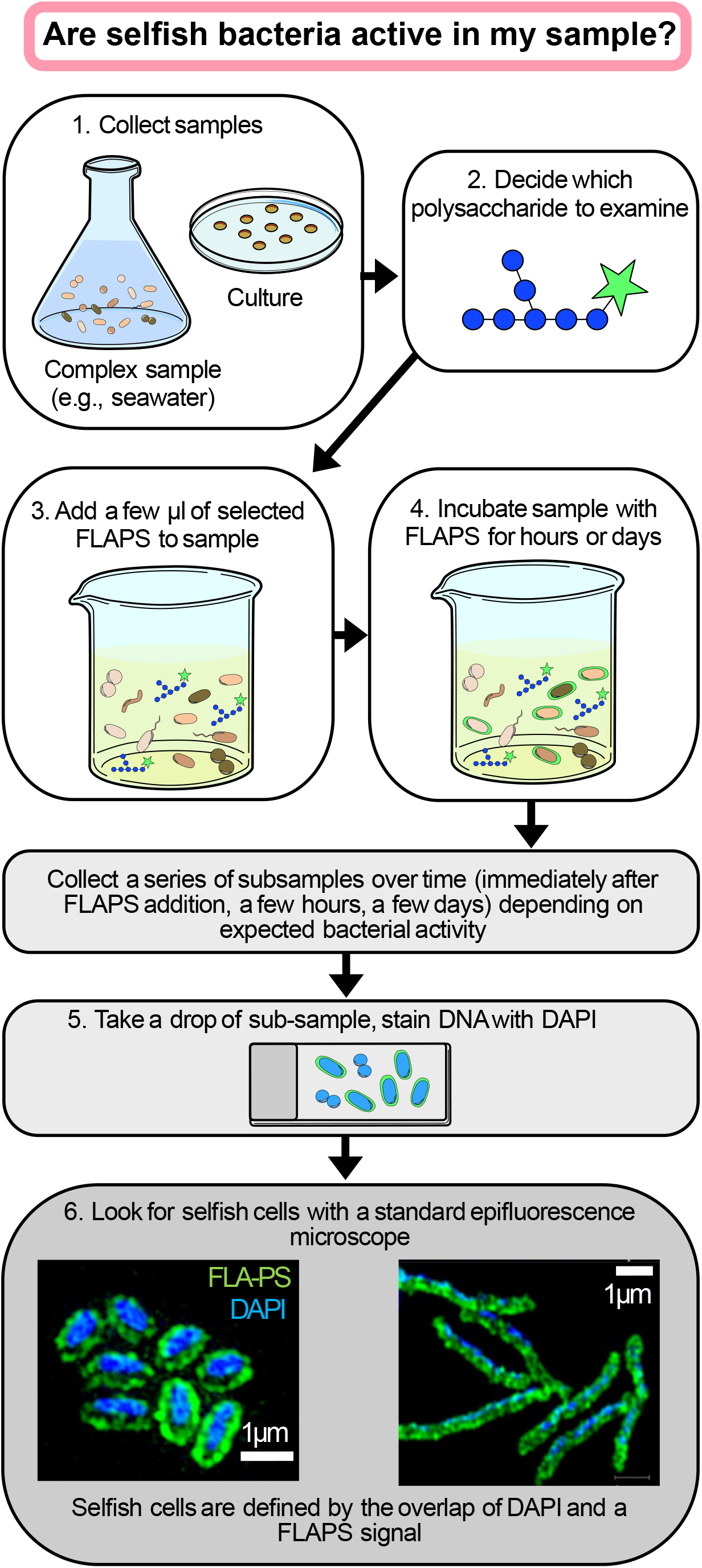
Simple workflow for the identification of selfish bacteria in pure cultures or complex samples through a fluorescently labeled polysaccharide (FLAPS) incubation experiment.

A wide range of soluble and semi-soluble polysaccharides can be labeled to probe a range of polysaccharide metabolisms (Supplementary Table S1, see Extended Methods for FLAPS production procedure). FLAPS have been successfully used with diverse pure cultures from different phyla and several environmental microbiomes, including marine (seawater and sediments ^23, 24, 27, 31^), limnic (unpublished), gut (unpublished), and rumen (cattle and sheep ^13,26,32^) (Supplementary Table S1).

### Measuring extracellular hydrolysis and selfish uptake of FLAPS – an environmental example

Using the procedures described above, we incubated an environmental sample - surface seawater collected in September off Helgoland (North Sea) - with three FLAPS (laminarin, xylan, and chondroitin sulfate) to determine the relative contributions of selfish bacteria and external hydrolyzers to polysaccharide degradation. External hydrolysis of polysaccharides, which produces low molecular weight hydrolysis products in the surrounding medium, was measured via gel permeation chromatographic analysis of the filtrate collected at each time point. Hydrolysis rates were calculated based on the shift in molecular weight classes as a polysaccharide is systematically hydrolyzed to lower molecular weight hydrolysis products over time (Fig. 1E).

The incubations showed rapid selfish uptake of both laminarin and xylan: 16% and 6% of cells were stained by these two FLAPS, respectively, already at the initial (t = 1 h) timepoint (Fig. 3 A). Selfish uptake increased up to 72 h; a low number of chondroitin-stained cells were also detected. All three polysaccharides were also externally hydrolyzed, with high hydrolysis rates of chondroitin and xylan detected in the 48 and 72 h samples, respectively (Fig. 3B). Total cell counts increased from 0.9 x10^9^ cells L^-1^ to 1.5 x10^9^ cells L^-1^ in all incubations (amended and unamended) within 24 h of incubation, but then diverged, with cell counts in the chondroitin incubations increasing to ca 3 x10^9^ cells L^-1^ at 72 h, a smaller increase in the xylan incubations, and a decrease in the laminarin and unamended incubations (Fig. 3 C). Counting total and selfish cells thus clearly answers the first key question: selfish bacteria were present and actively taking up all three FLAPS, but selfish uptake of laminarin and xylan was greater than for chondroitin.

**Fig. 3.**
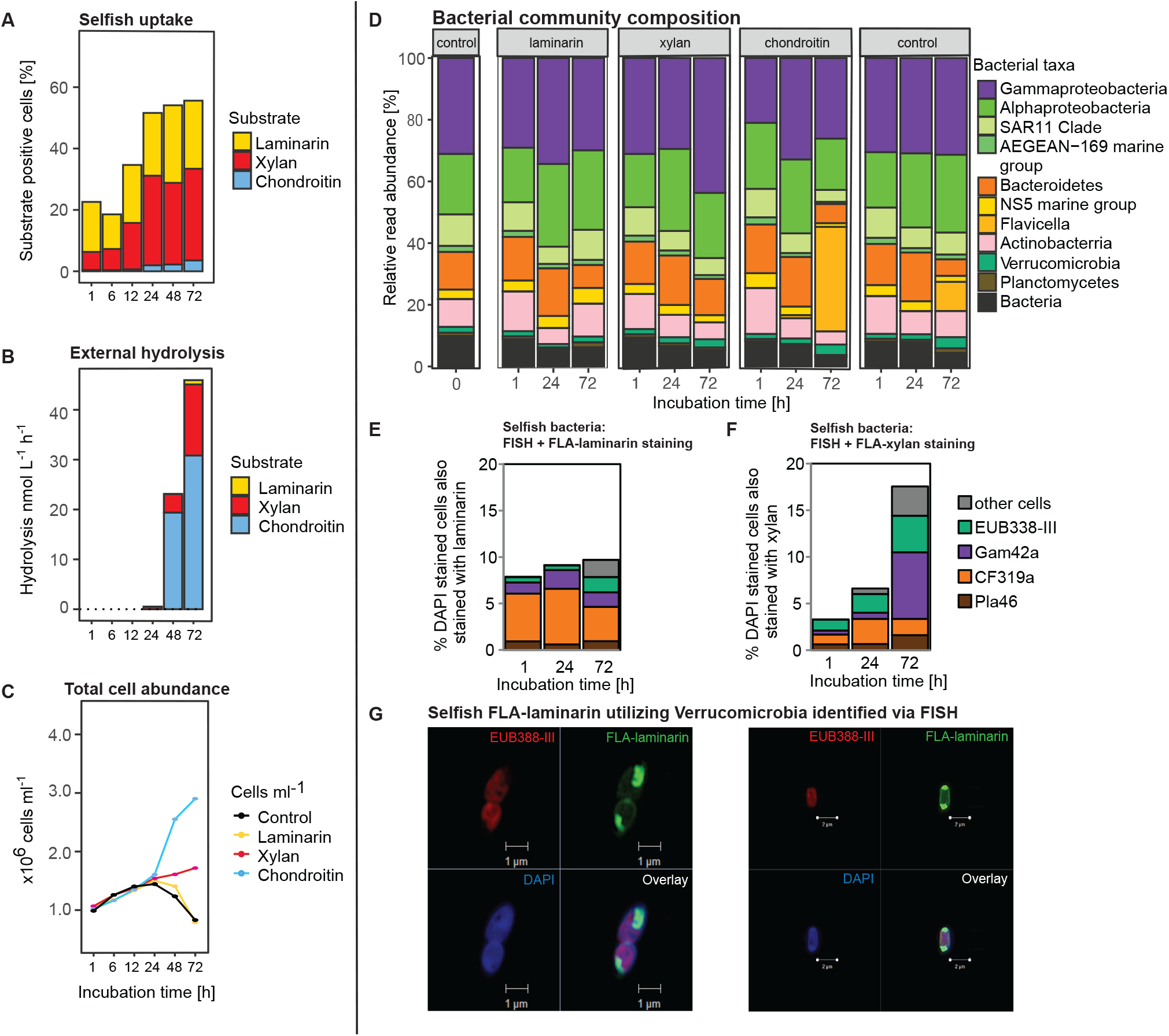
Polysaccharide utilization pattern in surface water from Helgoland in September over the course of a 72 h incubation. **(A)** Laminarin and xylan uptake stained a high proportion of cells already from the beginning of the incubation while **(B)** extracellular hydrolysis was comparatively rapid for xylan and chondroitin sulfate. **(C)** Microbial cell counts developed differently by substrate after the initial 24 h incubation. The **(D)** incubation-dependent changes of the initial community composition over 72 h show a FLA-chondroitin-dependent increase in *Flavicella* reads, but otherwise the community composition remained unchanged. Taxonomic identification with FISH showed **(E)** a large contribution of *Bacteroidetes* (CF319a) to selfish laminarin uptake while **(F**) a more diverse array of organisms including *Gammaproteobacteria* (Gam42a) and *Verrucomicrobia* (EUB338-III) took up xylan. **(G)** Super-resolution structured illumination images showing different polysaccharide accumulation pattern in FLA-laminarin stained cells (green), counterstained with DAPI (blue) and taxonomic correlation with FISH probes (red).

### Sequencing to gain insight into complex communities

After discovering whether selfish bacteria are present in a complex community, further questions may relate to bacterial identity: Which organisms are present in the initial sample? To what extent does the community change with increasing incubation time? Are there any indications of specific organisms responding to a FLAPS amendment? In these cases, amplicon sequencing of 16S rRNA genes (next-generation sequencing; NGS) can be the next step. In this case, unamended treatment controls (incubations to which no FLAPS were added) should be sequenced for the same time points as FLAPS-amended incubations to distinguish bottle effects from any substrate-dependent community responses. The correlation of a substrate-dependent change in bacterial taxa with a change in polysaccharide utilization can help identify potential taxa involved in the process^20^. Furthermore, selfish organisms can be phylogenetically stained and counted microscopically using fluorescence *in situ* hybridization (FISH^33^). Taxonomically specific FISH probes can be selected (or designed) to confirm the absolute abundance of a bacterial group and create a direct, visual link to selfish polysaccharide accumulation (Fig. 3 G).

The initial bacterial community in Helgoland waters in September was composed of *Gamma-* and *Alphaproteobacteria, Bacteroidetes*, and *Actinobacteria* (Fig. 3 D). Over the course of incubations, minor changes in abundance occurred within these groups. However, the bacteroidetal *Flavicella* was an exception, showing a large increase - of 34% and 10% - at 72 h in the chondroitin and control incubations, respectively, compared to the initial community.

### Revealing links between function and taxonomy – FISH on FLAPS-stained cells

Combining FLAPS uptake with FISH links an organism directly with its substrate, yielding information that is otherwise extremely difficult or impossible to obtain, particularly from environmental samples. Different FISH methods targeting rRNA can be used to visualize bacterial groups. Extensive testing (see Extended Methods) has demonstrated that modifying the protocol of Manz et al. (1992)^33^ using quadruple labeled oligonucleotide probes is most suitable for identifying FLAPS-stained selfish bacteria. It is compatible with FLAPS incubation because the procedure is less harsh when compared with other protocols (i.e., CARD-FISH see Extended Methods), has fewer steps, and gives a detectable FISH signal even for small cells from environmental samples.

Since the Helgoland incubations showed high selfish uptake of laminarin and xylan, we focused our FISH investigations on these samples, using probes targeting the abundant *Bacteroidetes* (CF319a) and *Gammaproteobacteria* (GAM42a) as well as the minor phyla *Verrucomicrobia* (EUB338-III) and *Planctomycetes* (PLA46), which have previously been found to be capable of selfish uptake ^34, 27, 29^. Laminarin incubations were clearly dominated by selfish *Bacteroidetes*, especially during the initial 24 h (Fig. 3 E), whereas in the xylan incubations, selfish *Gammaproteobacteria* and also *Verrucomicrobia* increased in proportion especially by 72 h of incubation (Fig. 3 F). We note, moreover, that the numbers of FISH- and FLAPS-positive cells are likely underestimated because the FISH procedure can lead to a loss of substrate signal in cells (Supplementary Fig. S2; see Extended Methods).

### Super-resolution microscopy – visualization of individual selfish substrate accumulation patterns

In addition to standard epifluorescence microscopy, high-resolution visualization of the accumulated FLAPS within the cell can be carried out using super-resolution structured illumination microscopy (SR-SIM) (Fig. 3G). Cells from the Helgoland FLA-laminarin incubation were identified as members of the *Verrucomicrobia* by FISH (Fig. 3G) and showed two different versions of polar substrate accumulation pattern with an enlarged periplasmic space, in contrast to elongated cells identified as *Gammaproteobacteria* that stained more evenly among the periplasm Fig. 3G). The use of a membrane stain can show the co-localization of the polysaccharide-associated green fluoresceinamine signal with the red membrane stain Nile Red in a fluorescent intensity line grating ^27^, further demonstrating polysaccharide uptake into the periplasmic space.

In addition, visualizing selfish uptake can reveal cellular heterogeneity in substrate processing (Fig. 4A). Bacterial cultures can exhibit homogenous or heterogenous staining patterns, as shown by two strains of *Bacteroidetes thetaiotaomicron* incubated with FLA-yeast mannan (Fig. 4A). Identifying and monitoring cell heterogeneity is essential in applications where cells are used for bioprocesses (e.g., production of fuels such as ethanol, butanol, fatty acid derivatives or natural products), as it affects biosynthesis performance, specifically enzyme activity or expression level ^35^. Mechanisms underlying microbial cell-to-cell heterogeneity that are not based on genotype are not well understood, but FLAPS incubation can visualize such heterogeneity.

**Fig. 4.**
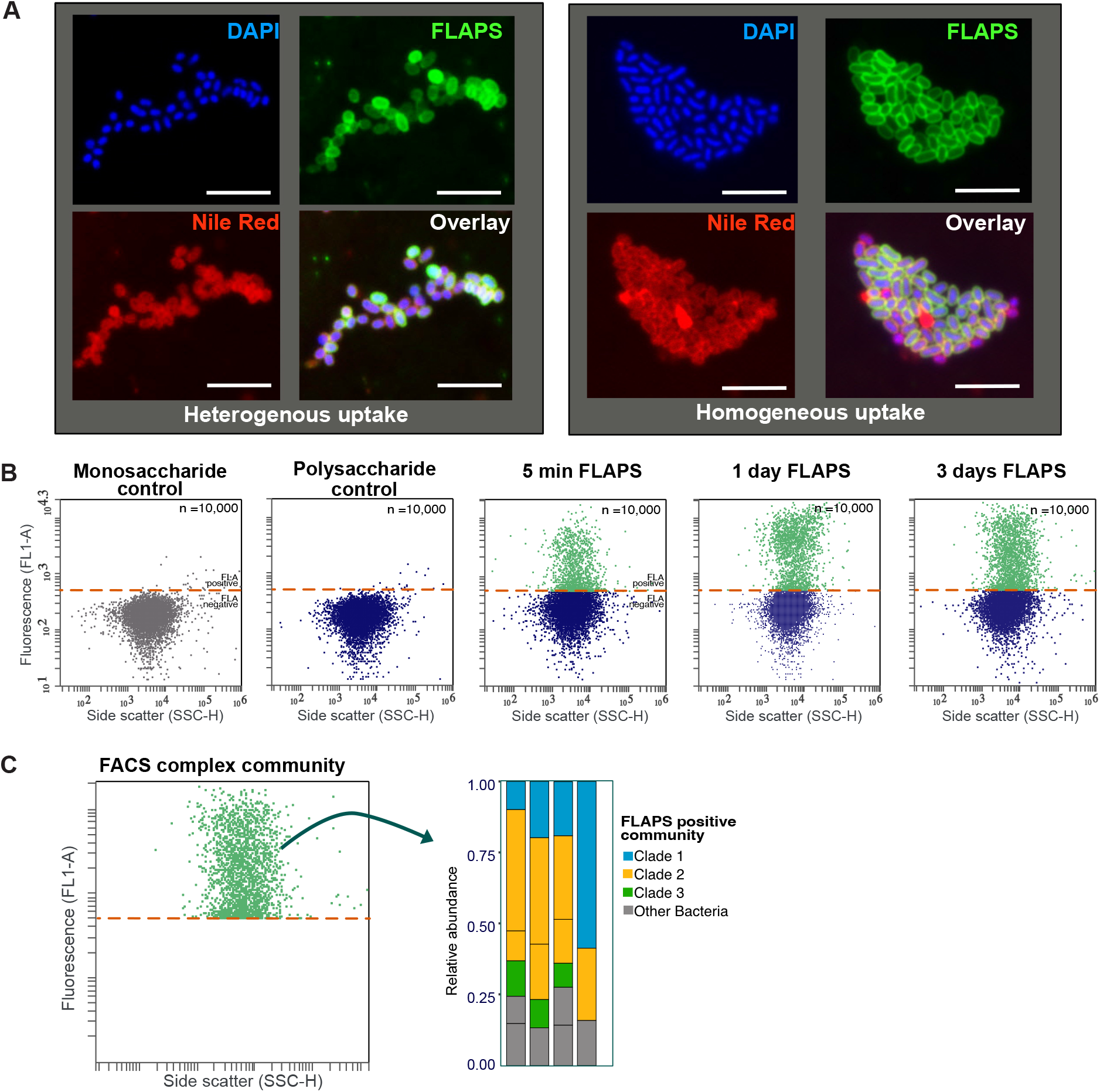
Metabolic phenotyping and flow cytometry of cultures and complex communities. **(A)** Visualization of cell-to-cell heterogeneity in strains of *Bacteroidetes thetaiotaomicron* incubated with fluorescently labeled yeast mannan. The cell DNA is shown by DAPI staining in blue, the FLAPS is shown in green and cell membranes are shown by Nile red staining in red. Scale bar = 5μm. **(B)** Quantification of FLAPS uptake into *Bacteroidetes thetaiotaomicron* by flow cytometry. Shown in the first two panels are negative controls: *Bacteroidetes thetaiotaomicron* grown on unlabeled monosaccharide and unlabeled polysaccharide. The subsequent three panels show the change in fluorescence of the cells with incubation in FLAPS for 5min, 1 day and 3 days. Data revisualized from Klassen et al., 2021. **(C)** Schematic representation fluorescence-activated cell sorting of FLAPS positive cells in combination with sequencing.

### Flow cytometry: tracking specific organisms

Especially for cases in which identification of specific cells is difficult, or – for pure cultures, for example – when the relative change in FLAPS uptake needs to be measured over short timescales, flow cytometry can be helpful. Bacteria that take up FLAPS can be flow cytometrically identified based on their physical and fluorescence properties within minutes and sorted based on the fluorescence signature of FLAPS accumulation (Fig. 4B, C). Flow cytometry can be used in environmental samples to identify subpopulations of FLAPS-stained cells. Various controls are required, including a blank control of the community with the nucleic acid counterstain for background noise calibration (Supplementary Fig. S3A). Since flow cytometry of killed controls is problematic (see Extended methods), a possibility for a negative control is the addition of FLAPS to a fixed and thus inactivated sample to account for any unspecific binding of the substrate (Supplementary Fig. S3C-I). For pure cultures, flow cytometry can be used to identify differences in uptake efficiency between cultures by plotting fluorescence intensity over forward scatter (proxy for cell size) or side scatter (proxy for cell granularity) (Fig. 4B) ^26^. The pure culture without FLAPS, as well as the medium without cells but with FLAPS, should be used to calibrate the background noise.

Fluorescence-activated cell sorting (FACS) of selfish populations can be used to assess the taxonomic composition and functional potential of active selfish organisms in a complex community. Here, selected bacterial populations are separated from the sample and enriched. Applying FISH on sorted cells can quantify taxa in a sorted population and link taxonomy to uptake pattern ^29,30^. Moreover, cells stained at different intensities with the FLAPS could be used as a proxy to differentiate between different populations within a sample. However, in environmental samples quantification of unstained vs. stained cells would only be a rough estimate, as there are potentially very pronounced differences in the staining pattern among different taxa at a given time (Fig. 3G). Note in all of these cases that microscopic validation of selected populations after sorting is necessary is advised. Additionally, a nucleic acid stain can be used as an independent parameter to ensure that cells (and not background signals) are sorted.

## Conclusions

To investigate the processing of polysaccharides by microorganisms and microbial communities, phenotypic approaches which allow for *in situ* probing are essential ^36^. Several major points emerge from our investigations of polysaccharide processing: most importantly, by overlooking selfish bacteria, a major substrate processing mechanism carried out by bacteria in a wide range of environments is missed. We note that external hydrolysis of laminarin was minimal in our incubations, yet selfish uptake was rapid, even in the initial community collected from the ocean (Fig. 3A-C). However, low selfish uptake of chondroitin shows that external hydrolysis can also be important – and that the importance of the polysaccharide processing mechanism varies by substrate, since the same starting communities were present in each incubation. Moreover, the ability to carry out selfish uptake is phylogenetically widespread: in the laminarin and xylan incubations, substantial selfish uptake did not correlate with changes of specific taxa in the bulk community analysis. This observation – and the broad range of selfish cells, especially in the xylan incubation at 72 h - suggests widespread prevalence of the selfish mechanism among diverse bacteria. Although selfish *Bacteroidetes* and *Verrucomicrobia* constitute a major portion of the total in early xylan incubations, *Gammaproteobacteria* constitute a large fraction of selfish bacteria at 72 h.

We emphasize here that selfish bacteria transport large polysaccharide fragments into the periplasmic space; they do not simply bind them to the outer membrane. Super-resolution light microscopy has shown that FLAPS staining is confined to the periplasmic space, which can be well-defined by the simultaneous use of a membrane stain ^27^. Fluorescence line profiling and z-stack images localizing the 3 dimensions of the cell demonstrate that the polysaccharide is within the outer membrane, but outside the cell wall ^27^. In addition to visual evidence, physiochemical proof of polysaccharide uptake is provided by work with mutant strains lacking the outer membrane uptake system for polysaccharides (SusC/D) or the full PULs. Unlike their wild-type counterparts, these mutant strains were not stained with FLAPS ^13, 26^. The mechanisms of selfish uptake of polysaccharides have been thoroughly studied among members of the *Bacteroidetes*, especially gut-associated strains. Studies have demonstrated that polysaccharides are bound at the outer membrane, partly hydrolyzed, and then transferred into the periplasmic space, where they are further hydrolyzed (Fig. 1 B) ^37-39,21^. Genes involved in selfish uptake by *Bacteroidetes* are typically possessed in PULS ^17, 42^, including the SusC/D transport system. Observations that selfish uptake is also carried out by organisms lacking PULs (e.g., *Planctomycetes*) and organisms lacking SusD (*Gammaproteobacteria*) demonstrate, however, that the presence of a PULs or a SusC/D system is not a strict requirement for selfish uptake. Moreover, bacteria with PULs and SusC/D systems may carry out external hydrolysis in addition to selfish uptake ^38, 39^. Examination of genomes without experimental incubations consequently is not sufficient to demonstrate selfish behavior. The specific means by which organisms lacking the SusC/D system carry out selfish uptake remain to be determined. In sum, broad use of a selfish strategy of substrate processing in the environment – partly by organisms whose specific mechanism of selfish uptake is not yet known – suggests that much is waiting to be revealed. The methods to do so are at hand.

## Online methods

### Sample collection and substrate incubation

Surface seawater was collected at the long-term ecological research station Helgoland Roads on the 17^th^ and 19^th^ September 2018. At both dates, FLAPS incubation experiments were conducted in sterile, acid-rinsed glass bottles in the dark at 18 °C (ambient water temperature). Incubations with FLA-laminarin, FLA-xylan, and FLA-chondroitin sulfate at a final concentration of 3.5 μM monomer equivalent were conducted in triplicates. Incubations without FLAPS addition and a single autoclaved (killed) control per substrate were also included. After 0, 1, 6, 12, 24, 48 and 72h of incubation, subsamples were taken for total cell counts, analysis of selfish uptake and external polysaccharide hydrolysis, FISH analysis and fluorescence-activated cell sorting. Subsamples for bacterial community analysis were taken at 0, 24, and 72 h. Quantitative data on phytoplankton community composition (cell counts of centric and pennate diatoms, dinoflagellates, coccolithophores and flagellates), and data on water temperature and salinity, nutrient availability (silicate, nitrate and phosphate), and Chl *a* concentrations were collected on the same dates ^29^.

### FLAPS production

FLAPS were produced according to Arnosti 2003^30^. Briefly, hydroxyl groups of a soluble polysaccharide are activated using cyanogen bromide. The activated polysaccharide is labeled by incubating with fluoresceinamine ((Fluoresceinamine, Isomer II, Sigma-Aldrich). Subsequently, the labeled polysaccharide is separated from unreacted fluoresceinamine and purified using size exclusion chromatography. See Extended Methods for a detailed description of FLAPS production and troubleshooting guide, as well as tips on labeling “problem” polysaccharides.

### Extracellular hydrolysis

External hydrolysis of FLAPS was measured by analyzing filtrate from the samples collected for bacterial community analysis (described below). These samples, collected after 0, 24, 48, and 72 h of incubation, were analyzed as described in detail in Arnosti (2003) ^30^. The earlier timepoints (6 and 12 h) were not measured because, in our experience, environmental samples from the water column do not show sufficient activity for hydrolysis to be detected at these early timepoints. Sediment incubations, and incubations with pure cultures of bacteria, in contrast, typically show much higher hydrolysis rates, and hydrolysis can be measured on timescales of hours ^26, 30^.

### Selfish uptake measurements

For all microscopic analysis, samples were fixed with formaldehyde at a final concentration of 1%, filtered onto a 0.2 μm pore size polycarbonate filter, counterstained with 4’,6-diamidino-2-phenylindole (DAPI), and mounted in a Citifluor/VectaShield (4:1) solution.

Total DAPI counts and selfish polysaccharide uptake were analyzed using a fully automated epifluorescence microscope (Zeiss AxioImager.Z2, Carl Zeiss), equipped with a Colibri LED light source (Carl Zeiss), a cooled charged-coupled-device (CCD) camera (AxioCam MRm, Carl Zeiss), and a HE-62 multifilter module (Carl Zeiss). A minimum of 45 fields of views were acquired, microscopic images exported into the modified image analysis software ACMEtool (M. Zeder, Technology GmbH, http://www.technobiology.ch and Max Planck Institute for Marine Microbiology, Bremen), and signals evaluated (Supplementary Table S2) according to Bennke *et al*. (2016)^40^. Images from the substrate channel were acquired at three exposure times (10 ms, 35 ms, and 140 ms) to cover the diversity of signal intensities and patterns of FLAPS accumulation. Automated counts were confirmed by manual microscopy; careful, manual curation of the processed images is a requirement. See Extended Methods for tips on dealing with background signals during microscopy.

For single-cell detection of substrate uptake patterns, cell membranes stained with Nile Red and FLA-substrate labeled cells were subsequently visualized using SR-SIM. Individual cells were analyzed with a Zeiss ELYRA PS.1 (Carl Zeiss) microscope equipped with 561, 488, and 405 nm lasers and BP 573-613, BP 502-538, and BP 420-480+LP 750 optical filters. A Plan-Apochromat 63x/1.4 Oil objective was used to take z-stack SR-SIM images with a CCD camera. Data processing and image analysis were done using the ZEN software package (Carl Zeiss).

### Flow cytometry

Single-cell fluorescence quantification was determined using an Accuri C6 flow cytometer (BD Accuri Cytometers, USA). The 8- and 6-peak validation beads (Spherotech, USA) were used for reference. All culture samples were measured under 488nm laser excitation, and the fluorescence was collected in the FL1 channel (530 ± 30 nm). The medium with and without fluorescent substrate and an electric threshold of 17,000 FSC-H were used to set the background noise. All bacterial samples, with and without FLAPS, were measured using a slow flow rate with a total of 10,000 events per sample in triplicate. Bacteria are detected from the signature plot of SSC-H vs. green fluorescence (FL1-H). The flow cytometric output was analyzed using the FlowJo v10-4-2 software (Tree Star, USA). See Extended Methods for details on flow cytometric and fluorescent-activated cell sorting parameters and troubleshooting guide.

### Bacterial community analysis and community statistics

In seawater incubation experiments, the initial bacterial community and changes in community composition and abundance were determined by 16S rRNA amplicon sequencing. At each time point, a subsample of 10 ml from each incubation bottle was filtered through a 0.2 μm pore size polycarbonate filter; the filtrate was used to analyze extracellular hydrolysis rates, as described above. Total DNA was extracted from the filter with the DNeasy Power Water Kit (Quiagen) and the hypervariable V3-V4 region (490 bp) of the 16S rRNA was amplified from the DNA using the S-D-Bact-0341-b-S-17 and S-D-Bact-0785-a-A-21 ^41^ primer pair with an Ion Torrent sequencing adapter and an Ion Xpress Barcode Adapter (Thermo-Fischer Scientific) attached to the forward primer. The PCR product was purified and remaining free primer were removed using the AMPure XP PCR Cleanup system (Beckman Coulter). A pool of barcoded PCR products in equimolar concentration was further amplified in an emulsion PCR with the Ion Torrent One-Touch System (Thermo Fischer Scientific). Sequencing was done on an IonTorrent PGM™ sequencer (Thermo Fischer Scientific) in combination with the High-Q™ View chemistry (Thermo Fischer Scientific). Quality trimmed (> 300 bp sequence length, < 2% homopolymers, < 2% ambiguities) reads were demultiplexed and used as input for the SILVAngs pipeline ^42^ for taxonomic assignment of the reads based on sequence comparison to the SSU rRNA SILVA database 312.

### Fluorescence *in situ* hybridization for taxonomic identification

Combining FISH with FLAPS incubations allows the correlation of taxonomy and function, as defined by the capability of bacteria for selfish polysaccharide uptake.

We tested the effect of two FISH procedures on FLAPS signals. For this, we took two seawater samples from the sampling station Kabeltonne off the island of Helgoland and incubated them for 48 h with a) FLA-laminarin and b) FLA-xylan. Subsequently, we fixed the samples with formaldehyde at a final concentration of 1% for 1 h. We applied a tetra-labeled FISH^33^ and the CARD-FISH^43^ protocol to the incubations to test if FISH influences the microscopic evaluation of FLAPS-stained cells (see Extended Methods for details). For both procedures, we used probes for the taxonomic identification of most Bacteria, *Planctomycetes*, and *Verrucomicrobia* (EUB388-I, PLA46, and EUB388-III, respectively, Supplementary Table S3). Formamide concentrations in the hybridization buffer were probe-specific (Supplementary Table S3). The number of FLAPS-stained cells after FISH treatment, the co-localization of the FLAPS and FISH signal, and the taxonomic correlation of FLAPS-labeled cells were evaluated using epifluorescence microscopy combined with automated image analysis.

Both FISH procedures cause wash-out of FLAPS signal from the cells, which was dependent on the harshness and number of steps in the procedure (Supplementary Fig. S2 & S4). For more details, see the extended methods section, where we give a detailed description of the comparison and also included a guide of how to handle FLAPS signal background. We recommend the use of the tetra-labeled FISH procedure to minimize signal loss.

Based on the results of the methodological comparison, the abundance of FLAPS-stained *Gammaproteobacteria, Bacteroidetes, Verrucomicrobia*, and *Planctomycetes* was analyzed on samples from Helgoland FLA-laminarin and FLA-xylan incubations using 4 x Atto594 labeled probes (GAM42a, CF319a, EUB388-III + competitor EUB338-II, and PLA46, respectively, Supplementary Table S3).

## Acknowledgements

We thank our colleagues from Biologische Anstalt Helgoland, especially Antje Wichels, Eva-Maria Brodte, and Uwe Nettelmann, for enabling the sampling campaign on Helgoland and facilitating substrate incubations and sample preparation. In addition, we thank the crew of the *Aade* for sample collection, Maria Belen Gonzalez Pino for help with incubation experiments, sample preparation, and microscopic analysis, and Sherif Ghobrial (UNC) for help with the work on the fluorescently labeled polysaccharides. GR has received funding from the European Union’s Horizon 2020 research and innovation program under the Marie Sklodowska-Curie grant agreement No. 840804 and the Deutsche Forschungsgemeinschaft (DFG, German Research Foundation) Project number 496343779. This study was supported and funded by the Max Planck Society and the U.S. National Science Foundation (OCE-1736772 and -2022952 to CA).

## Author contributions

All authors conceived different aspects of the experimental study; GG performed sample collection, GG conducted FLAPS incubations; GG and GR performed FISH, flow cytometry, and microscopy analyses; CA synthesized FLAPS and analyzed samples for external hydrolysis; GG, CA, and GR analyzed the data; RA, CA. GR secured funding; GG, GR and CA produced the figures and tables; all authors contributed to writing the manuscript.

