## Supplementary Figure for "Visualizing and identifying selfish bacteria: a hunting guide"

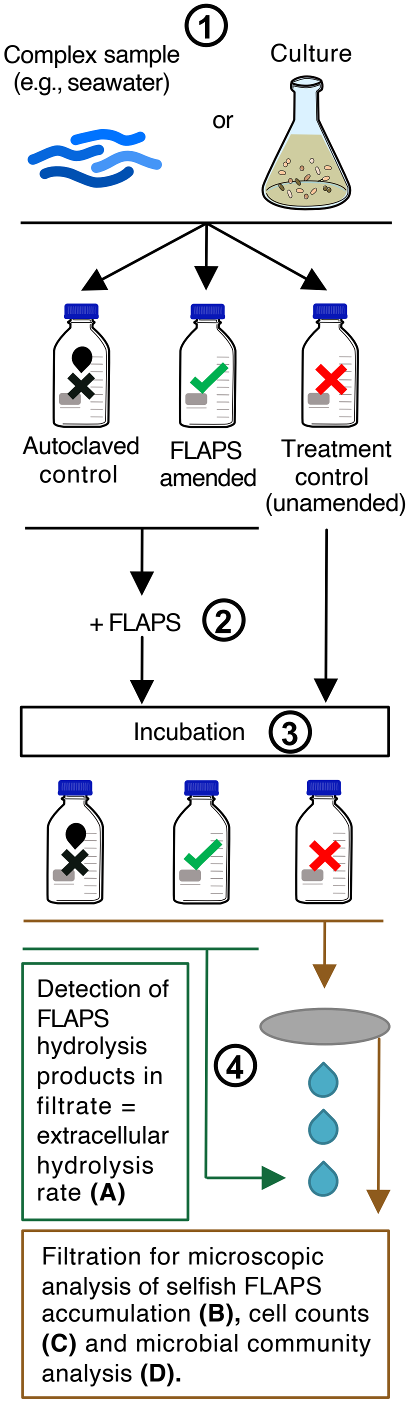


**Supplementary Figure S1** Workflow to analyze extracellular polysaccharide hydrolysis **(A)**, selfish polysaccharide uptake **(B)**, total cell counts **(C)** pure cultures and the underlying microbial community **(D)** from complex samples. **1** Priming with unlabeled polysaccharides of pure cultures show the general potential to utilize polysaccharides **2** Low or ambient concentrations testing for the natural potential to utilize polysaccharides. High concentrations to show the general potential to utilize polysaccharides. **3** Long incubation (days - weeks) testing for the natural potential to utilize polysaccharides in natural waters. Short incubation (hours - days) to show the general potential to utilize polysaccharides or the natural potential to use polysaccharides in sediments or other cell-rich natural environments. **4** Fixation of the sample might be required for further analysis. Formaldehyde fixation, for example, stabilizes the cell structure, stabilizes the substrate in the cell and maintains morphology for microscopic analysis.

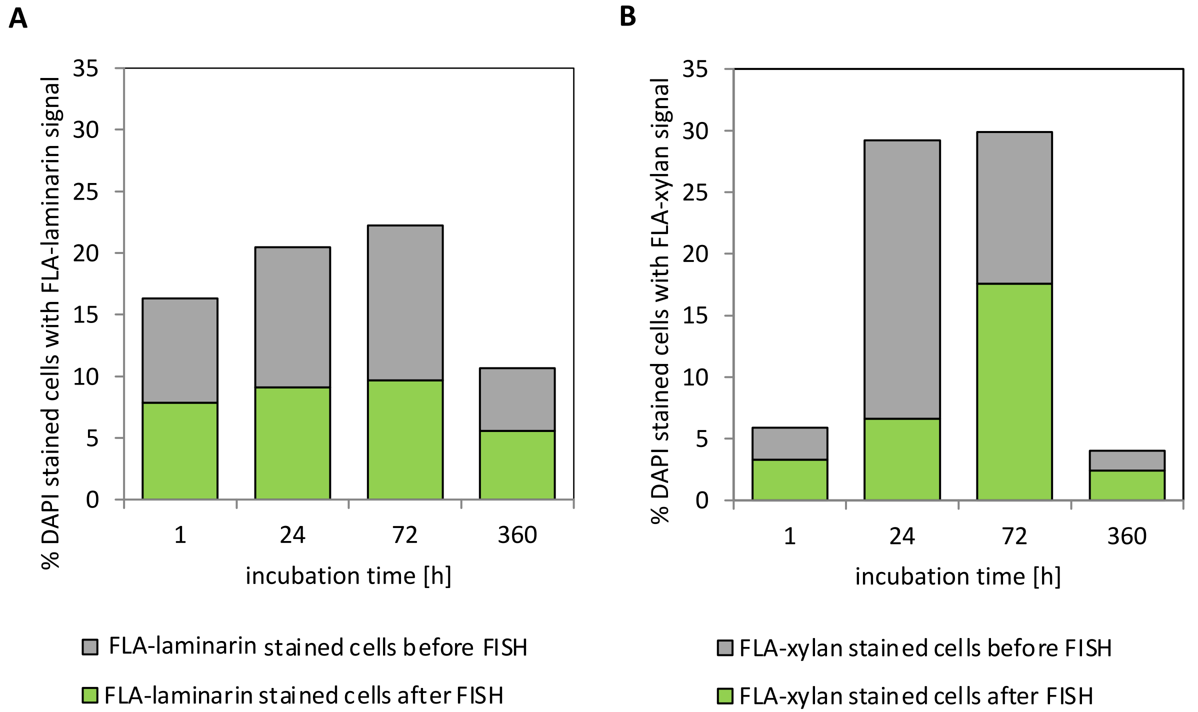


**Supplementary Figure S2** Proportion of **(A)** FLA-laminarin stained cells and **(B)** FLA-xylan stained cells before and after FISH treatment.

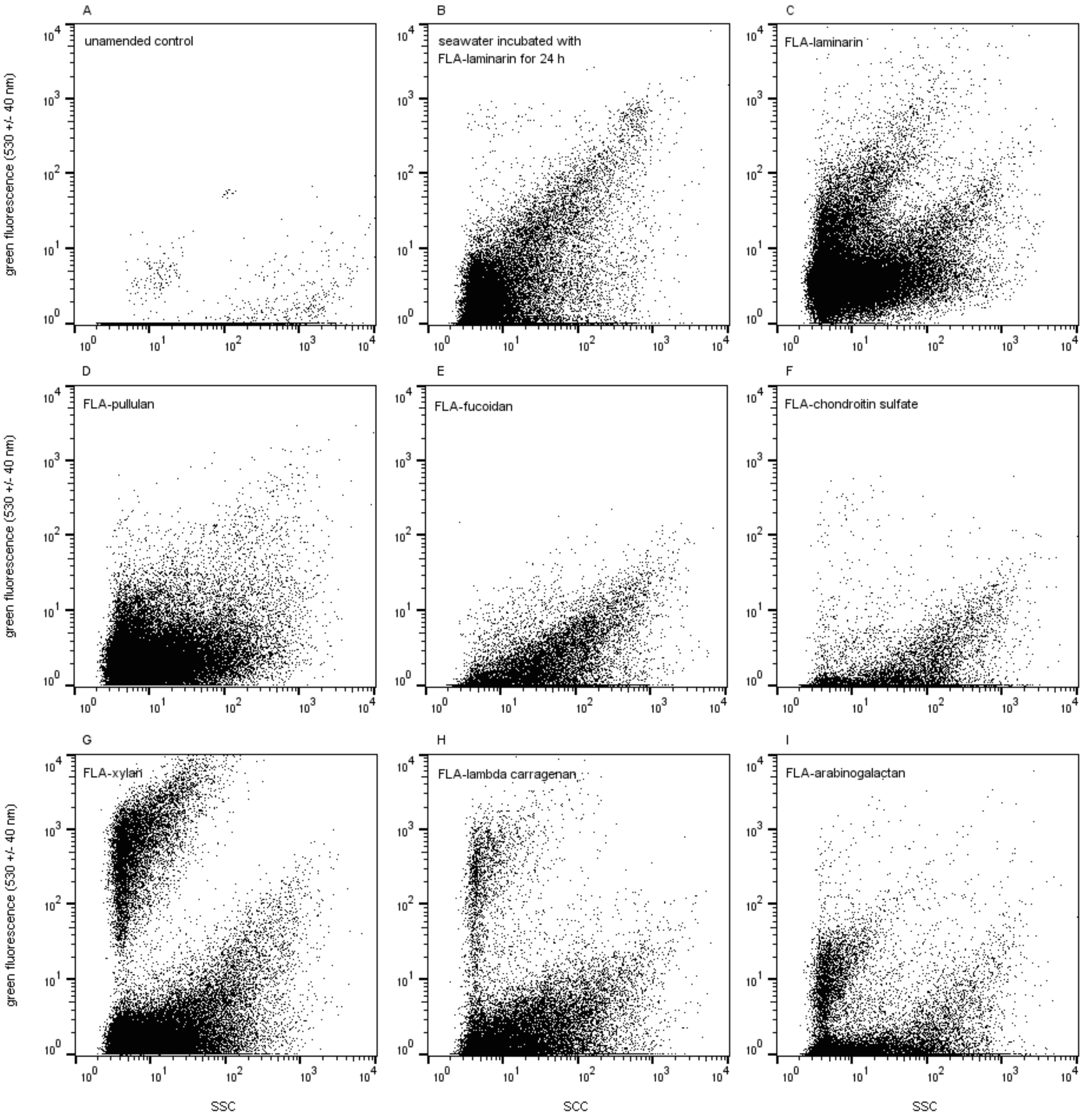


**Supplementary Figure S3** Flow cytometric dot plots showing FLAPS background signals in marine surface waters. **(A)** Seawater without the addition of any FLAPS to define the samples background noise. **(B)** Seawater sample containing some FLA-laminarin stained cells after 24 h of incubation. Formamide fixed seawater supplemented with **(C)** FLA-laminarin, **(D)** FLA-pullulan, **(E)** FLA-fucoidan, **(F)** FLA-chondroitin sulfate, **(G)** FLA-xylan, **(H)** FLA-lambda carrageenan and **(I)** FLA-arabinogalactan to define substrate background noise.

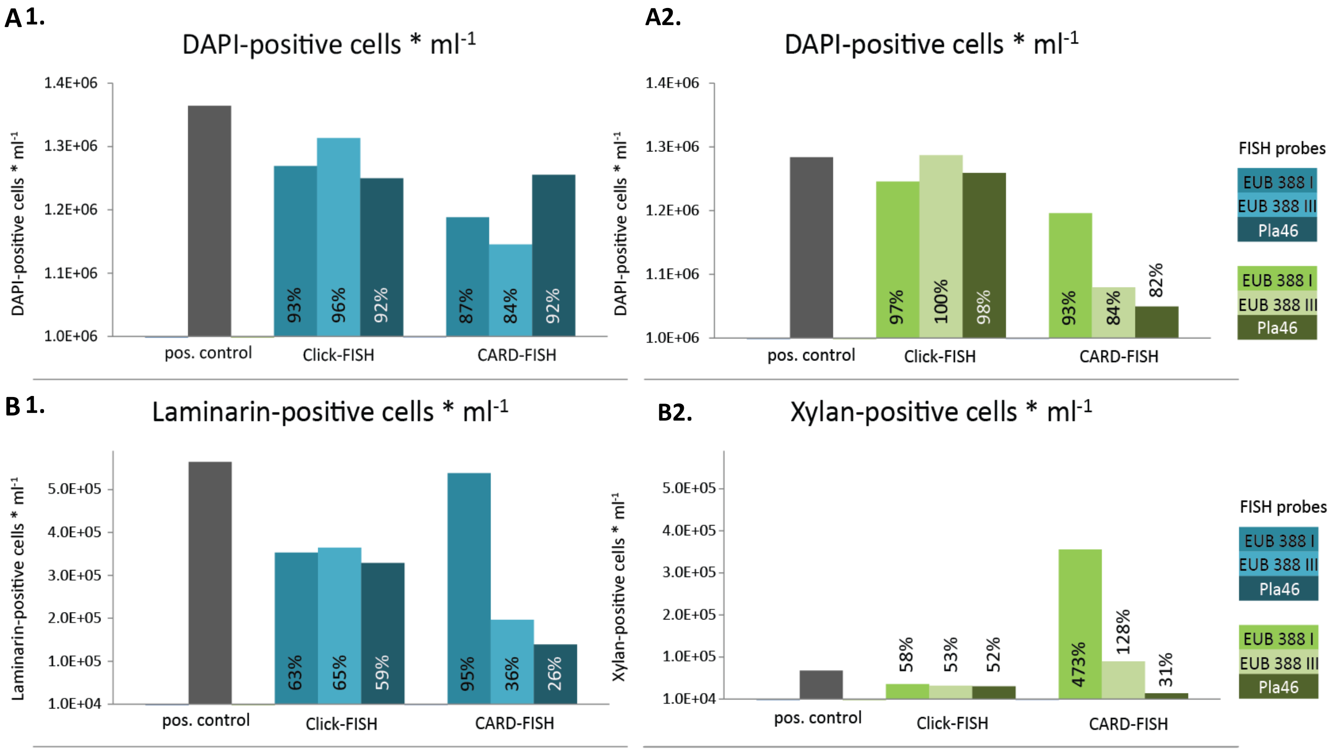


**Supplementary Figure S4** Comparison of FLA-substrate labeled cells from summer incubations of Helgoland seawater, taxonomically stained using tetra-labeled FISH probes and CARD-FISH; **(A1&2)** DAPI-positive cells without treatment (positive control (pos. control)) and after FISH treatment in the 1) laminarin and 2) xylan incubations. The percentages of DAPI-positive counts after FISH treatment relates to total DAPI-positive cells in pos. control. **(B1&2)** Substrate-positive cells without treatment (pos. control) and after FISH treatment in the 1) laminarin and 2) xylan incubations. The percentages of substrate-positive counts after FISH treatment relates to total substrate-positive cells in pos. control

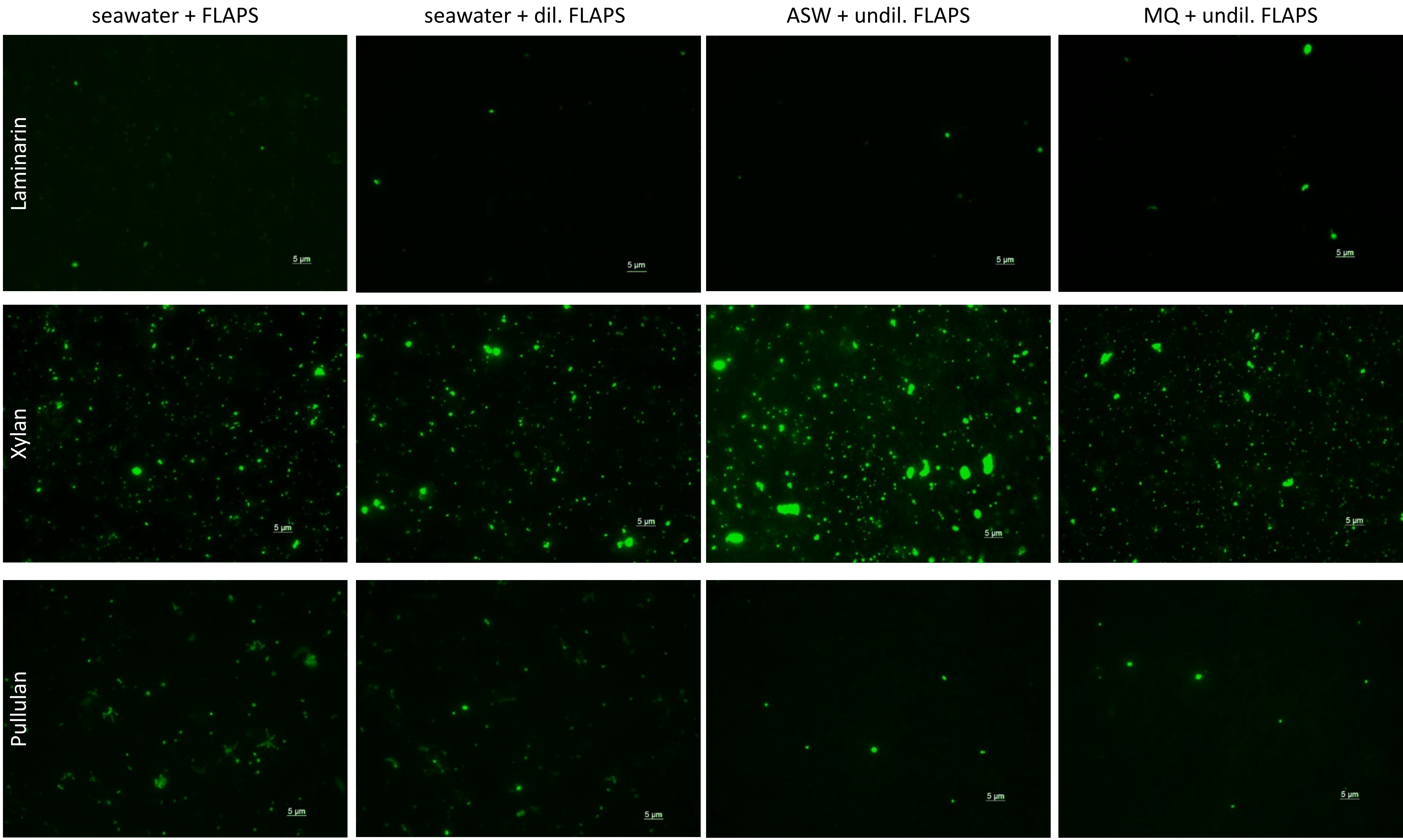


**Supplementary Figure S5** Comparison of FLAPS and DAPI staining is required to identify substrate-stained cells. The background noise of fluorescently labeled polysaccharides (FLA-PS) after substrate dilution in different solvents. ASW = artificial seawater, MQ = 18 MΩcm water.

| Polysaccharide | Sugar Type | Monosaccharide units | Habitat | Bacterial culture | Extracellular hydroylsis (EE) /Selfish | Reference |
|---|---|---|---|---|---|---|
| **Alginic acid** | 1,4-glycosidic bond | β-D-mannuronate and α-L-guluronate | Seawater, oxic and anoxic marine sediment | | EE | Arnosti, 2003 |
| **Arabinogalactan** | β-1,3, β-1,5, β-1,6 (Galactan), α-1,3, α-1,5, (Arabinan) | Arabinose and galactose | Seawater (Coastal, temperate and gyre regions) | *Gramella forsetti* | EE / Selfish | Reintjes et al., 2017 & 2020, Balmonte et al., 2018 & 2019 |
| | | | Oxic and anoxic marine sediment | | EE | Arnosti & Holmer, 2003 |
| | | | Freshwater (limnic) | | EE / Selfish | unpublished |
| **Amylopectin** | α-1,4 | glucose | | *Bacteroidetes thetaiotaomicron* | Selfish | unpublished |
| **Chondrotin Sulfate** | β-1,3, β-1,4 | N-acetylgalactosamine, glucoronic acid | Seawater (Coastal, temperate and gyre regions), marine particles | | EE / Selfish | Reintjes et al., 2017 & 2020, Balmonte et al., 2018 & 2019, Manna et al., 2022 |
| | | | Oxic and anoxic marine sediment | | EE | Arnsoti, 2003, Arnosti & Jørgensen 2006 |
| | | | Freshwater (limnic) | | | unpublished |
| **Direct microalgae extract** | Diverse | Diverse | | | | |
| *Isochrysis* | | | Seawater, mesocosum | | | Murry et al., 2007 |
| *Skeletonema sp.* | | | Seawater, mesocosum | | | Murry et al., 2007 |
| *Spirulina* | | | Seawater mesocosum | | | Arnosti 2008 |
| *Wakame* | | | Seawater mesocosum | | | Arnosti 2008 |
| **Direct macroalgae extract** | Diverse | Diverse | | | | |
| *Mazzaella japonica* | | | Anaerobic rumen and artifical rumen system | *Bacteroidetes thetaiotaomicron* | Selfish | unpublished |
| *Saccharina latissima* | | | Anaerobic rumen and artifical rumen system | *Bacteroidetes thetaiotaomicron* | Selfish | unpublished |
| *Alaria esculenta* | | | Anaerobic rumen and artifical rumen system | *Bacteroidetes thetaiotaomicron* | Selfish | unpublished |
| *Macrocystis pyrifera* | | | Anaerobic rumen and artifical rumen system | *Bacteroidetes thetaiotaomicron* | Selfish | unpublished |
| *Asparagopsis taxiformis* | | | Anaerobic rumen and artifical rumen system | *Bacteroidetes thetaiotaomicron* | Selfish | unpublished |
| *Ulva sp.* | | | Anaerobic rumen and artifical rumen system | *Bacteroidetes thetaiotaomicron* | Selfish | unpublished |
| *Ascophyllum nodosum* | | | Anaerobic rumen and artifical rumen system | *Bacteroidetes thetaiotaomicron* | Selfish | unpublished |
| **Fucoidan** | Diverse | fucose (galactose, xylose, arabinose, rhamnose) | Seawater (Coastal, temperate and gyre regions) | | EE / Selfish | Reintjes et al., 2017 & 2020, Balmonte et al., 2018 & 2019 |
| | | | Oxic and anoxic marine sediment | | EE | Arnsoti, 2003, Arnosti & Jørgensen 2006 |
| | | | Freshwater (limnic) | | Selfish / EE | unpublished |
| **Glycogen** | α-1,4. α-1,6 | Glucose | | *Bacteroidetes thetaiotaomicron* | Selfish | unpublished |
| **Galactan** | α-1,3, α-1,6 | Galactose | | *Bacteroidetes thetaiotaomicron* | Selfish | unpublished |
| **Inulin** | β-2,1 | Fructose | | *Bacteroidetes thetaiotaomicron, Bacteroidetes ovatus* | Selfish | unpublished |
| **Laminarin** | β-1,3, β-1,6 | Glucose | Seawater (Coastal, temperate and gyre regions), marine particles | *Gramella forsetti* | EE / Selfish | Reintjes et al., 2017 & 2020, Balmonte et al., 2018 & 2019, Manna et al., 2022 |
| | | | Oxic and anoxic marine sediment | | EE | Arnsoti, 2003, Arnosti & Jørgensen 2006 |
| **Levan** | β-2,6 | Fructose | | *Bacteroidetes thetaiotaomicron, Bacteroidetes ovatus* | Selfish / EE | unpublished |
| **Mucin** | Diverse | Diverse | Seawater | | Selfish / EE | unpublished |
| | | | Marine sediment | | Selfish / EE | unpublished |
| **Porphyran** | β-1,3, α-1,4 | galactosyl, galactosyl 6-sulfate or 3,6-anhydrogalactosyl | | *Bacteroides plebeius, Bacteroides uniformis, Bacteroidetes thetaiotaomicron, Bacteroidetes ovatus* | Selfish | Robb et al., 2022 |
| **Pullulan** | α-1,4, α-1,6 | Glucose | Seawater (Coastal, temperate and gyre regions), marine particles | *Gramella forsetti* | Selfish / EE | Reintjes et al., 2017 & 2020, Balmonte et al., 2018 & 2019, Manna et al., 2022 |
| | | | Oxic and anoxic marine sediment | | EE | Arnsoti, 2003, Arnosti & Jørgensen 2006 |
| **Rhamnogalacturonan II** | diverse | Homogalacturan, galacturonic acid, galactose, rhamnose, arabinose, galactan | | *Bacteroidetes thetaiotaomicron* | Selfish | Hehemann et al., 2019 |
| **Starch** | α-1,4 | Glucose | | *Bacteroidetes thetaiotaomicron* | Selfish | unpublished |
| **Xylan** | β-1,4, β-1,3 | Xylose | Seawater (Coastal, temperate and gyre regions), marine particles | | Selfish / EE | Reintjes et al., 2017 & 2020, Balmonte et al., 2018 & 2019, Manna et al., 2022 |
| | | | Oxic and anoxic marine sediment | | EE | Arnsoti, 2003, Arnosti & Jørgensen 2006 |
| | | | Freshwater (limnic) | | Selfish / EE | unpublished |
| **Yeast-mannan** | α-1,6 (backbone), α-1,3, α-1,2 (side chains) | Mannose | Anaerobic rumen and mouse gastrointestinal tract | *Bacteroidetes thetaiotaomicron* | Selfish | Hehemann et al., 2019, Klassen et al., 2010 |
| **λ-carrageenan** | β-1,3, α-1,4 | galactopyranose, 3,6-anhydrogalactopyranose | Seawater (Coastal, temperate and gyre regions) | | Selfish / EE | Reintjes et al., 2020 |
| | | | Anaerobic rumen | *Bacteroidetes thetaiotaomicron* | Selfish | unpublished |

**Supplementary Table S1** Polysaccharides that have been fluorescently labelled for the extracellular enzyme (EE) or selfish uptake analysis. Highlighted are the habitats and organisms in which these polysaccharides have been tested. Note that studies carried out prior to 2012 did not in any case include investigation of selfish uptake. See Arnosti et al. 2011 for a partial summary of prior investigations in marine planktonic environments.

**Supplementary Table S2** Recommended settings for the evaluation of FLA-polysaccharide stained cells and their taxonomic identification by fluorescence *in situ* hybridization with an automated epifluorescence microscope and the image analysis software ACMETool; 1 pixel = 0.10601 $\mu m^2$. SBR = signal-to-background ration; MGV = medium grey value.

| Dye | Excitation wavelength [nm] | Exposure time [ms] | Signal definition (ACMETool) | | |
|---|---|---|---|---|---|
| | | | Area [pixel] | SBR | MGV |
| 4′,6-Diamidin-2-phenylindol (DNA stain, UV) | 365 ± 4.5 | 25 | 18 – 150 | 2 | 55 |
| Fluoresceinamine (FLA-polysaccharide) | 470 ± 14 | 140<br>35<br>10 | 18 – 200<br>18 – 250<br>18 – 200 | 1.8<br>2<br>2 | 55<br>65<br>55 |
| 4xAtto594 (Mono-FISH) | 590 ± 17.5 | 110 | 18 – 400 | 2 | 55 |
| Atto594 (CARD-FISH) | 590 ± 17.5 | 70 | 18 – 400 | 2 | 55 |
| Auto fluorescence | 590 ± 17.5 | 500 | >6 | | |

**Supplementary Table 3** Fluorescence *in situ* hybridization probe overview.

| Probe | Target organisms | Sequence (5' -> 3') | Formamide [%] | Reference |
|---|---|---|---|---|
| EUB338-I | Bacteria | GCTGCCTCCCGTAGGAGT | 35 | Amann *et al.,* 1990 |
| EUB338-III | *Verrucomicrobia* | GCTGCCACCCGTAGGTGT | 35 | Daims *et al.,* 1999 |
| PLA46 | *Planctomycetes* | GACTTGCATGCCTAATCC | 30 | Neef *et al.,* 1998 |
| CF319a | *Bacteroidetes* | TGGTCCGTGTCTCAGTAC | 35 | Manz *et al.,* 1996 |
| GAM42a | *Gammaproteobacteria* | GCCTTCCCACATCGTTT | 35 | Manz *et al.,* 1992 |

**Supplementary Table S4** Treatment overview for substrate background signal testing. FOV = field of view;ASW = artifical seawater; MQ = 18 MΩcm water; FA = formaldehyde; FLAPS = fluorescently labeled polysaccharide; EDTA = Ethylenediaminetetraacetic acid; - = not tested; N.D. = not detectible due to overexposure.

| | | Background signal counts at 35 ms/ FOV | | | | | |
|---|---|---|---|---|---|---|---|
| | Treatment: FLAPS + [X] | Laminarin | Xylan | Chondroitin sulfate | Pullulan | Arabino-glactan | Fucoidan |
| A | X = seawater + 1% FA | 23 | 1553 | 9 | 316 | 12 | 2 |
| B | X = ASW + 1% FA | 166 | N.D. | 82 | N.D. | 145 | 216 |
| C | X = MQ + 1% FA | N.D. | N.D. | 725 | N.D. | 527 | N.D. |
| D | X = ASW | 18 | 1078 | 3 | 86 | 8 | 2 |
| E | X = MQ | 17 | 1130 | 10 | 720 | 5 | 3 |
| F | X = seawater + 1% FA, FLAPS = 100 x dilluted | 42 | 1112 | 5 | 204 | 10 | 1 |
| G | X = seawater + 25 mMol EDTA | N.D. | N.D. | N.D. | - | - | - |
| H | X = ASW + 25 mMol EDTA | N.D. | N.D. | N.D. | - | - | - |
| I | X = MQ + 25 mMol EDTA | 91 | 1195 | 143 | - | - | - |
| J | X = ASW + 5 min at 32 °C prewarmed FLAPS | 26 | 1630 | N.D. | - | - | - |
| K | X = ASW + 15 min at 32 °C prewarmed FLAPS | 321 | 1403 | 48 | - | - | - |
| L | X = MQ + 5 min at 32 °C prewarmed FLAPS | 104 | 1847 | 446 | - | - | - |
| M | X = MQ + 15 min at 32 °C prewarmed FLAPS | 93 | 1591 | 499 | - | - | - |
| N | X = seawater + 5 min at 32 °C prewarmed FLAPS | 405 | 1743 | N.D. | - | - | - |
| O | X = seawater + 15 min at 32 °C prewarmed FLAPS | 215 | 1525 | N.D. | - | - | - |
