## Extended Methods for "Visualizing and identifying selfish bacteria: a hunting guide"

*[#]shared first authorship*

### Correspondence

*Greta Reintjes, University of Bremen, Bibliothekstraße 1, 28359, Bremen, Germany, mail:, phone: +49 421 2028 9710

### Keywords

selfish uptake; carbon cycling; enzymatic hydrolysis; bacterial community function; flow sorting; polysaccharide degradation

### Running title

Identifying selfish bacteria

### Supplementary methods

#### Synthesis and characterization of fluorescently labeled polysaccharides (FLAPS)

Many polysaccharides can be used for FLAPS experiments (Supplementary Table S1). FLAPS are prepared using highly purified polysaccharides obtained from commercial sources or direct extracts purified to a specific molecular weight cutoff. The polysaccharides are activated using cyanogen bromide (CNBr), which reacts with hydroxyl groups of the polysaccharide to form cyanate esters. These groups react readily with primary amines. The activated polysaccharide is then incubated with the fluorophore fluoresceinamine (Fluoresceinamine, Isomer II, Sigma-Aldrich). During labeling, the primary amine group on the fluoresceinamine linker forms a stable isourea linkage with the activated polysaccharide. Although other primary amine-containing fluorophores can, in principle, be used to label polysaccharides, we note that fluorophores with an ester group in their linker arm are unsuitable for experiments in aqueous solution because the ester group is very easily auto-hydrolyzed (Arnosti, unpublished data). After activation, the labeled polysaccharide is separated from the unreacted fluorescent tag and purified and characterized using size exclusion chromatography or centrifugation with membrane cartridges and multiple rounds of washing (Arnosti, 1995 & 2003; Glabe *et al.,* 1983).

In principle, any soluble polysaccharide (or phytoplankton extract or phytoplankton-derived DOC; Arnosti, 2003; Arnosti *et al.,* 2005; Murray *et al.,* 2007) can be labeled using this procedure. The primary limitations on polysaccharide choice are the requirements for solubility in an aqueous solution and stability (and solubility) at pH > 9.5. The labeling density of FLAPS is highly polysaccharide dependent (likely related to the hydrodynamic volume and conformation of a polysaccharide, and thus to access of a fluorophore to activated polysaccharide sites). It must be determined for each batch of FLAPS. Labeling with a fluorescent tag is not thought to change the bioactivity of a polymeric compound (Glabe *et al.,* 1983).

#### Extracellular hydrolysis measurements

The rate of extracellular FLAPS hydrolysis is determined by the change in size fractions from the initial polysaccharides to smaller hydrolysis products, as determined via gel permeation chromatography, as described in detail in Arnosti*,* 2003.

#### Fluorescence *in situ* hybridization on FLAPS-stained cells.

In this study, we systematically compared the effect of performing tetra-labeled FISH and CARD-FISH on FLAPS-stained cells. Tetra-labeled FISH (referred to as just FISH) is fast, consisting of one hybridization and two washing steps. However, the four fluorophores per probe restrict the signal intensity, and therefore the probe concentration must be high to saturate all possible binding sites (0.84 µMol ~5 ng DNA $\mu l^{-1}$). Comparatively, the CARD-FISH protocol includes more steps: embedding, cell permeabilization, inactivation, and a CARD-reaction step. The CARD reaction activates numerous fluorochromes per probe - enhancing signal intensity compared to directly labeled probes - at a lower probe concentration (0.028 µM or 0.16 ng DNA $\mu l^{-1}$). Ethanol washing steps must be left out of both protocols as ethanol's permeabilization of the cell wall was found to lead to FLAPS signal loss.

For these experiments, we took two seawater samples from the sampling station Kabeltonne off the island of Helgoland, Germany. We incubated them for 48 h with FLA-laminarin and FLA-xylan. Subsequently, we fixed the samples with formaldehyde at a final concentration of 1% for 1 h at room temperature. The cells were then filtered onto 0,2 µm pore size polycarbonate filters using a gentle vacuum of < 200 mbar. We applied FISH and CARD-FISH with the probes for the taxonomic identification of most Bacteria, *Planctomycetes*, and *Verrucomicrobia* (EUB388-1, PLA46, and EUBI388-III, respectively, Supplementary Table S3). Formamide concentrations in the hybridization buffers were probe-specific (Supplementary Table S3). After the FISH treatments, all samples were counterstained with 4',6-diamidino-2-phenylindole (DAPI) and mounted in a Citifluor/VectaShield (4:1) solution. We co-localized the DAPI (DNA) FLAPS and FISH signal and evaluated the cellular abundance using epifluorescence microscopy with automated image acquisition and enumeration software (Bennke et al., 2016).

Firstly, we found that the FISH and CARD-FISH treatments caused 4 ± 3% and 13 ± 4% loss of the total cell signals (DAPI), respectively (Supplementary Figure S4 A1&2). Furthermore, FISH caused a loss of 48 ± 3% and 52 ± 3% of the FLAPS signal (laminarin- and xylan-positive signals, respectively, Supplementary Figure S4 B). Comparatively, a minimum of 71 ± 2% FLAPS signals were lost after the CARD-FISH protocol. (Supplementary Figure S4 B1&2).

Due to the high FLAPS signal loss during CARD-FISH, we recommend using a tetra-labeled probe with the FISH protocol after Manz *et al.,* (1992) to identify FLAPS-stained cells. It should be noted that even with FISH, there is a FLAPS signal loss and that the numbers of FISH- and FLAPS-positive cells are likely underestimated (Supplementary Figure S3). Optimizations of the FISH protocol to reduce signal loss should be performed.

Furthermore, FISH staining in the far-red spectrum (594 nm) can cause crosstalk that facilitates false-positive detection of FISH signals as substrate signals (488 nm). We recommend an emission filter with a reduced bandwidth for the green spectrum.

**Flow cytometry and fluorescence-activated cell sorting (FACS)**

For flow cytometric analysis from seawater incubations, cells were fixed with formaldehyde (1% final concentration), and counterstained with DAPI to differentiate between substrate background signal and substrate-stained cells. Cells were flow cytometrically measured using a BD Influx™ Cell Sorter (Becton-Dickinson, Germany), equipped with a 100.0 mW UV-laser (Coherent, Germany) with an excitation wavelength of 355 nm and a 200.0 mW Coherent Saphire laser (Coherent, Germany) with an excitation wavelength of 488 nm. Sheath fluid contained 0.15% NaCl, and the nozzle has a diameter of 70 µm. Forward scatter (FSC), DAPI, and green fluorescence polysaccharide signal were detected by the photomultiplier tubes (PMT) 1 via nm band-pass filter at 14.0 V, PMT 3 via 530 ± 40 nm band-pass filter at 55.0 V and PMT 7 via 460 ± 50 nm band-pass filter at 45.0 V, respectively. Fluorescbrite® Multifluorescent Microspheres 1.0 µm (Polysciences, Inc., U.S.) were used as a standard for the alignment of the flow cytometer, triggering on FSC. Data were recorded with the BD FACS™ Software version 1.2 (BD lifescience, Germany) and analyzed with the FlowJo® v10 flow cytometry analysis software (Tree Star, U.S.).

Moreover, it is possible to perform cell sorting of selected populations. Therefore, the sorter is to be aligned using, e.g., 6 µm FACS™ Accudrop Beads (BD lifescience, Germany), and sorting should be performed in the 1.0 Drop Single mode (Giljan *et al.,* 2023). About 10 000 cells should be sorted into low-binding collection tubes for further microscopic analysis.

### Technical issues with respect to FLAPS synthesis and analysis

#### Preparation of 'problem' polysaccharides and polysaccharide-containing extracts or concentrates

Polysaccharides that are sparingly soluble (such as xylan) can be sonicated and filtered prior to activation (see Arnosti, 2003); gel-forming polysaccharides such as lambda-carrageenan and mucin can be solubilized in very dilute solution (e.g., a few mg $mL^{-1}$, compared to 20 mg $mL^{-1}$ for highly soluble polysaccharides) and labeled. Changing the pH of the initial solution may also increase solubility.

Initial polysaccharide preparation may also be necessary for polydisperse polysaccharides or concentrates, or extracts. Since extracellular enzymatic activity is detected as the change in molecular weight (MW) class of an added polysaccharide, the initial polysaccharide should ideally be high MW, covering a narrow MW range rather than polydisperse. Many commercially available polysaccharides (including fucoidan and chondroitin), as well as extracts and concentrates, are highly polydisperse. During the synthesis procedure, the MW range can be observed if the activated polysaccharide runs through a UV/Vis detector as it elutes (measuring absorption at 290 nm is often sufficient even in the case of neutral polysaccharides, provided the detector is reasonably sensitive). If the initial polysaccharide solution is highly concentrated (e.g., 20 mg $mL^{-1}$), a narrow molecular weight range can be collected by simply following the UV/vis detector signal and collecting - real-time - just the initial part of the polysaccharide peak in the activation vial. If the initial polysaccharide is too polydisperse, the polysaccharide can either be dialyzed (e.g., using 5000 or 10000 MWCO dialysis membranes), or the HMW fraction can be concentrated using spin cartridges with 5000 or 10000 MWCO limits. Dialyzed solutions must subsequently be lyophilized to obtain sufficient starting material; concentrates from spin cartridges should be repeatedly washed and concentrated to remove low MW fractions.

Polysaccharide-containing extracts can also be obtained from phytoplankton, as described in detail in Arnosti *et al.* (2005). In this case, cells were concentrated by centrifugation, ground, and extracted with organic solvents (acetone; chloroform: methanol) to remove lipids, and the resulting cellular mass was refluxed in an aqueous solution with 0.01% SDS. Following dialysis and lyophilization, carbohydrate content can be characterized, and the carbohydrate-containing soluble fraction can be labeled as described above and in Arnosti (2003) and as applied in Arnosti (2008).

### FLAPS background signals and how to handle them

#### Differentiating between signal and noise

Even though the application of the FLAPS substrate incubations is straightforward, differentiating between a substrate-stained cell and substrate background noise requires practice. The accumulation of FLAPS in the cells' periplasm results in a regular outline of the cells' morphology; therefore, irregular shapes hint at the presence of background noise. Nevertheless, different patterns in FLAPS accumulation within the cells lead to variations in the distribution and intensity of the FLAPS signal (Giljan *et al.,* 2023). Total cell counts should be determined for each incubation time point to test if a drop-in selfish activity correlates with an increase in cell number. Additionally, the distribution of substrate to daughter cells might lead to fluorescence dilution within a cell after division.

Flow cytometry can also be used to detect selfish uptake. However, as the detection time is in the microsecond range (Muratori *et al.,* 2008), sensitivity is lower than in a good microscope, and therefore strong staining of the target cell compared to background signals is necessary.

#### Background signal test of the FLAPS and how to solve it

The application of FLAPS in seawater or salt-containing medium can result in the production of non-cell associated background signals (Supplementary figure 3). These unspecific signals are caused by the spontaneous aggregation of the FLAPS to gel-like particles through contact with bivalent cations, leading to a size transition from dissolved to particulate organic matter. Furthermore, FLAPS can also adhere to, e.g., *in situ* particulate organic matter, creating background signals. These unspecific signals must be differentiated from true signals to achieve accurate quantification of FLAPS uptake.

Three polysaccharides that frequently show unspecific substrate signals (xylan, laminarin, and pullulan) were used to test background signal reduction. We tested several alterations to the standard FLAPS protocol (Supplementary Table S4). All FLAPS were mixed with 10 ml (A) seawater from sampling station Kabeltonne off the island Helgoland, (B) 1 x sterile filtered artificial seawater (ASW), and (C) 18 MΩcm water (MQ-water) at a concentration of 3.5 µM monomer equivalent final concentration. After inoculation, all solutions were supplemented with 1% Formaldehyde (FA) and fixed at room temperature for 1h. An unfixed non-reference was prepared for (D) ASW and (E) MQ-water. To check if the direct contact of the highly concentrated polymers with bivalent cations catalyzes a polymerization process of the substrates, the FLAPS were diluted 100-fold with MQ before the addition to (F) Helgoland water. For FLA-xylan, FLA-laminarin, and FLA-pullulan, ethylenediaminetetraacetic acid (EDTA) as an anti-chelating agent was added at a final concentration of 25 nM to the seawater (G), ASW (H) and MQ-water (I) to compete for the $Ca^{2+}$Ions and reduce polymerization. To avoid the addition of already polymerized substrates, FLAPS solutions were warmed to 32 °C for 5 or 15 minutes before addition to ASW (J, K), MQ (L, M), or seawater (N, O), respectively (Supplementary Table S4). Each 10 ml sample was filtered onto a 0.2 µm pore size polycarbonate filter, and microscopic pictures were taken for automated image analysis as described in detail in the methods section.

The substrate diluted in seawater with subsequent FA fixation (Supplementary Figure S5 left panel; Supplementary Table S4 A) was taken as the reference for a standard incubation in the marine environment. When laminarin, xylan, and pullulan were diluted in sterile artificial seawater, all background signal from unspecific binding to organic matter from an environmental sample was removed for pullulan but not for laminarin and xylan. To test whether bivalent cations from the seawater initialize the polymerization process, the FLAPS were added to sterile filtered MQ water. We found that the dilution of FLAPS in ultrapure water was a major cause of increased background signals if formaldehyde was added in addition, but the addition of formaldehyde did not appear to cause additional background signals in seawater. Neither the filtration of the substrate stock to remove particles larger than 0.22 µm from the substrate stock before the addition to the incubation nor the addition of EDTA as an anti-chelating agent led to a decrease in substrate background signal. EDTA addition led to the even distribution of the FLA substrate across the whole filter and led to complete overexposure (marked with N.D. in Supplementary Table S4).

Additionally, we tested the pre-warming of the substrate stock to dissolve potential aggregates of polysaccharides. Pre-warming of the substrate stock for 5 or 15 minutes to 32 °C did not remove the background signal. The background signal slightly decreased in number after 15 minutes at 32 °C for laminarin and xylan.

For background signal visualization in a flow cytometer, 2 ml 1% FA fixed seawater from the Helgoland autumn sampling was each mixed with the fluorescently labeled laminarin, xylan, and, pullulan, at a final concentration of 3.5 µM monomer equivalent and analyzed in a BD Influx$^{TM}$ Cell Sorter (Supplementary Figure S3 C-I). The substrate-specific background signal can be seen in comparison to an unamended treatment control (Supplementary Figure S3 A) and used as a negative control to separate it from FLAPS-stained cells. However, the comparison of the same sample, incubated with FLA-laminarin for 24 h showed that indeed the microbial community changes the substrate signature over the time of the incubation (Supplementary Figure 3 B+C).

### References from extended methods and supplementary figures
